# Catabolism of extracellular glutathione supplies amino acids to support tumor growth

**DOI:** 10.1101/2024.10.10.617667

**Authors:** Fabio Hecht, Marco Zocchi, Emily T. Tuttle, Nathan P. Ward, Bradley Smith, Yun Pyo Kang, Juliana Cazarin, Zamira G. Soares, Mete Emir Ozgurses, Huiping Zhao, Colin Sheehan, Fatemeh Alimohammadi, Lila D. Munger, Dhvani Trivedi, Gloria Asantewaa, Sara K. Blick-Nitko, Jason J. Zoeller, Ying Chen, Vasilis Vasiliou, Bradley M. Turner, Alexander Muir, Jonathan L. Coloff, Joshua Munger, Gina M. DeNicola, Isaac S. Harris

## Abstract

Restricting amino acids from tumors is an emerging therapeutic strategy with significant promise. While typically considered an intracellular antioxidant with tumor-promoting capabilities, glutathione (GSH) is a tripeptide of cysteine, glutamate, and glycine that can be catabolized, yielding amino acids. The extent to which GSH-derived amino acids are essential to cancers is unclear. Here, we find that GSH catabolism promotes tumor growth. We show that depletion of intracellular GSH does not perturb tumor growth, and extracellular GSH is highly abundant in the tumor microenvironment, highlighting the potential importance of GSH outside of tumors. We find supplementation with GSH can rescue cancer cell survival and growth in cystine-deficient conditions, and this rescue is dependent on the catabolic activity of γ-glutamyltransferases (GGTs). Finally, pharmacologic targeting of GGTs’ activity prevents the breakdown of circulating GSH, lowers tumor cysteine levels, and slows tumor growth. Our findings indicate a non-canonical role for GSH in supporting tumors by acting as a reservoir of amino acids. Depriving tumors of extracellular GSH or inhibiting its breakdown is potentially a therapeutically tractable approach for patients with cancer. Further, these findings change our view of GSH and how amino acids, including cysteine, are supplied to cells.

## Introduction

Amino acids are crucial to cancer initiation, progression, and drug resistance^1–3^. Despite their indispensability, amino acids are often scarce within the tumor microenvironment^4–6^, driving tumors to enable strategies to import and synthesize them^7–12^. Thus, interfering with these mechanisms and restricting amino acids holds significant potential as an anti-cancer strategy^13–16^.

One amino acid that has garnered significant attention as an anti-cancer target is cysteine^17,18^. Beyond being a building block for proteins, cysteine has other roles, including supporting the generation of antioxidants (e.g., glutathione, persulfide species)^19^, metabolites (e.g., H_2_S, CoA, hypotaurine)^20–23^, and iron-sulfur clusters for mitochondria^24^. The intracellular cysteine pool is suggested to be maintained by at least three sources: import through alanine/serine/cysteine/threonine transporter 1 (ASCT1), uptake of cystine through system x_c_^-^ (xCT and CD98) and subsequent reduction by thioredoxin reductase (TXNRD1), and de novo synthesis from methionine through the transsulfuration pathway. Cystine uptake through system x_c_^-^ is suggested to be the predominant source of cysteine in cancer cells^25^. Interestingly, the deletion of *Slc7a11* (which encodes xCT) in animals results in viable offspring^26^, suggesting that tissues (and potentially tumors) can obtain cysteine from another source. The generation of cysteine from the transsulfuration pathway is a potential source; however, this pathway is reported to be inactive in most tissues, and in the few tissues in which it is active, its activity is reduced in tumors from the corresponding tissue^27^. Together, this suggests that tumors potentially have an alternative mechanism to acquire cysteine.

Glutathione (GSH) is an antioxidant that regulates oxidative stress, drug detoxification, and post-translational modifications^28–30^. Disrupting GSH production is shown to impair tumorigenesis^31–33^, but the exact mechanisms involved are unclear. GSH is a tripeptide made of glutamate, cysteine, and glycine. Interestingly, GSH can be broken down into its individual constituent amino acids. The rate-limiting step in extracellular GSH catabolism is controlled by γ-glutamyltransferases (GGTs), which cleave the γ-glutamyl bond in GSH, releasing glutamate and dipeptide cysteinylglycine. After its release, cysteinylglycine is further broken down by peptidases to produce cysteine and glycine. GSH catabolism is proposed as a source of amino acids for cells^34–36^. This evidence dates back to seminal work from Harry Eagle, where supplementation with extracellular GSH was suggested to support cell growth in cystine-free conditions^37^. Indeed, mitochondrial GSH can be catabolized to supply cysteine for iron-sulfur cluster synthesis^38^. Unlike the deletion of *Slc7a11* in animals, deletion of the GGT family member *Ggt1* in animals results in higher GSH levels in the urine, lower cysteine levels of tissues, and perinatal lethality^39^. Notably, patients with mutations in *GGT1* often present with developmental disorders along with accumulation of GSH (i.e., glutathionemia, glutathionuria) and lower circulating levels of cystine^40–46^. Together, these findings suggest that GSH catabolism by GGTs is potentially an important source of amino acids, including cysteine. Whether tumors can co-opt GSH catabolism to fuel their growth is poorly understood.

Here, we identify an underappreciated role for GSH as a cysteine reservoir for tumors. We find that, compared to cystine, GSH can be highly abundant in the microenvironment of tumors. We show that supplementation with GSH or its product cysteinylglycine rescues cancer cells in cystine-depleted conditions. Mechanistically, we find that GGT activity is necessary and sufficient to promote survival by catabolizing GSH and supplying amino acids to surrounding cells. Further, we demonstrate that supplementation with extracellular GSH renders cancer cells resistant to drugs that block cystine uptake or its reduction into cysteine. Finally, we find that inhibiting GGT activity deprives tumors of cysteine and slows their growth. These results reveal an actionable pathway of nutrient acquisition in cancer with direct therapeutic implications. Further, these findings change our perspectives surrounding GSH biology and the acquisition of amino acids by cells.

## Results

### Production of GSH in tumors is dispensable for their growth

GSH facilitates tumorigenesis; however, it was unclear whether the intracellular production of GSH by tumors themselves was required. To test this, we bred the MMTV-PyMT transgenic mouse strain (which spontaneously develops breast tumors)^47^ with an inducible whole-body knockout mouse strain for the rate-limiting step in GSH synthesis, glutamate-cysteine ligase catalytic subunit (GCLC) (Rosa26-CreERT2 Gclc^f/f^)^48,49^. After allowing tumors to develop in these mice, the tumors were excised and orthotopically implanted into recipient immunocompetent wild-type mice (i.e., C57BL/6). The recipient mice were treated with tamoxifen, activating Cre recombinase activity and deleting *Gclc* specifically in the implanted tumors (Fig. 1a). *Gclc* mRNA and GSH levels were reduced in tumor-specific *Gclc* KO mice (Fig. 1b-1c), validating the model. Surprisingly, deletion of *Gclc* in tumors failed to perturb their growth (Fig. 1d-1e), suggesting that the tumor’s ability to synthesize intracellular GSH was dispensable. The lack of effect also suggested an alternative hypothesis where extracellular GSH was critical to tumor growth. To explore this, we examined GSH levels in the plasma and tumor interstitial fluid (TIF)^50,51^, the extracellular compartment surrounding tumors, from murine breast tumors (4T1 orthotopic allograft tumors). Analysis of plasma and TIF showed abundant total GSH levels, which were higher than reported levels from cell culture medium formulations (Fig. 1f). Further, total GSH levels were elevated in TIF compared to plasma. In line with these findings, analysis of pancreatic ductal adenocarcinoma (PDAC) from the *LSL-Kras*^G12D/+^ *Trp53*^f/f^ *Pdx-1-Cre* (KP-/-C)^5^ showed total GSH levels to be elevated in TIF in comparison to plasma and cell culture media (Extended Data Fig. 1a). These findings suggest that while intracellular GSH does not impact tumor growth and survival, extracellular GSH is highly abundant and potentially supports tumors.

**Figure 1.**
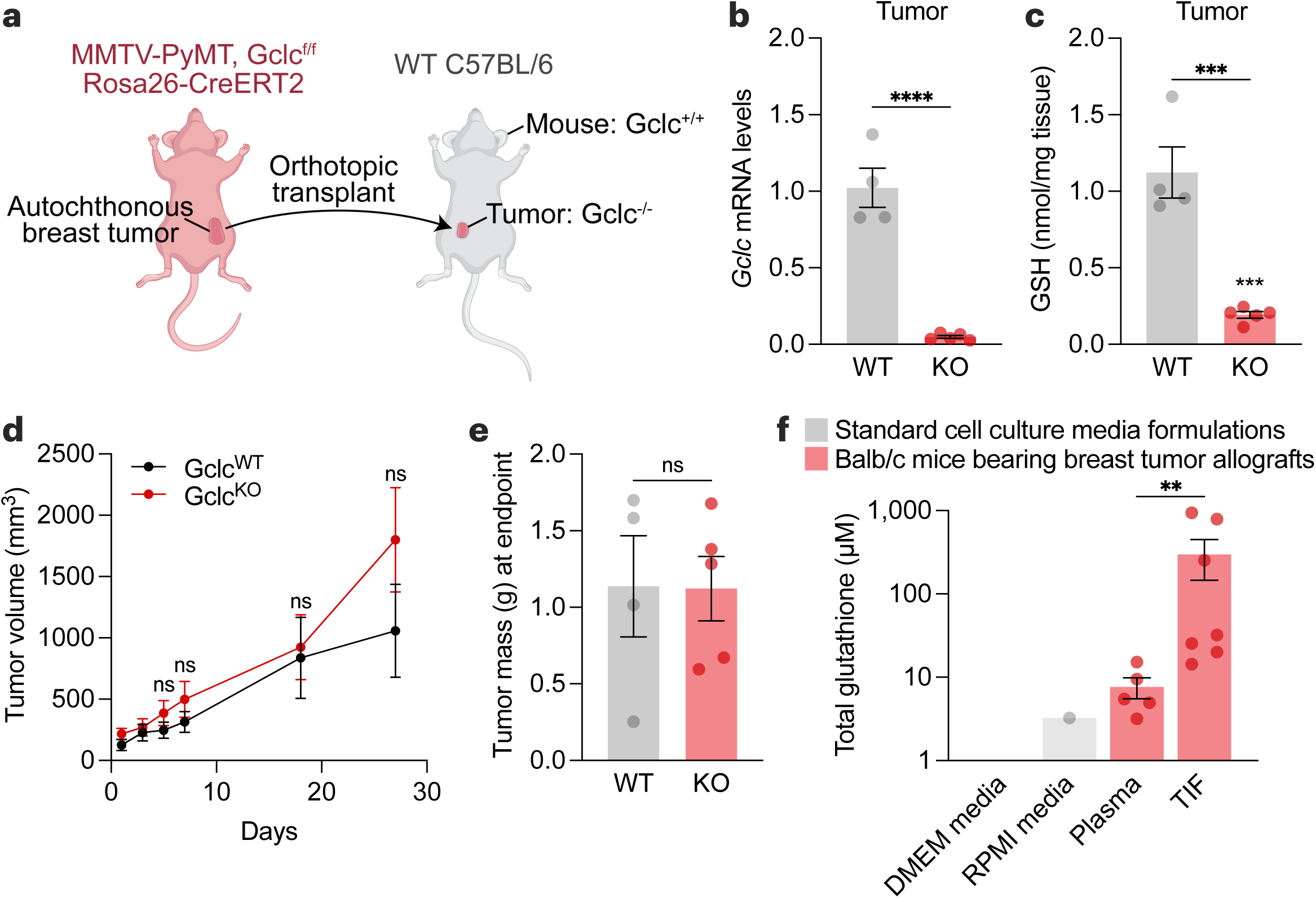
Intracellular production of GSH is dispensable for tumor growth. **a**, Schematic of the tumor-specific *Gclc* knockout mouse model. Autochthonous tumors from MMTV-PyMT Gclc^f/f^ Rosa26-CreERT2 mice were excised and orthotopically transplanted into mammary fat pads of wild-type C57BL/6 mice. C57BL/6 mice were treated with vehicle (corn oil; WT) or 160 mg/kg tamoxifen for 5 days (KO). **b-c**, Relative Gclc mRNA levels **(b)** and GSH levels **(c)** of WT tumors (n=4) and KO tumors (n=5). Statistical significance assessed by unpaired two-tailed t test. **d-e**, Tumor volume **(d)** and mass **(e)** of mice described in **(a-c)**. Statistical significance was assessed by two-way ANOVA (p=0.38) followed by Šídák’s multiple comparisons test for tumor volume and by unpaired t test (p=0.97) for tumor mass. **f**, Concentration of total glutathione in cell culture media formulations (grey bars), and in tumor interstitial fluid (n=7) and matched plasma (n=5) from Balb/c mice bearing 4T1 orthotopic allografts (red bars). Statistical significance was assessed by Mann-Whitney test. Data represented as mean ± s.e.m., ns, not significant; *p-value<0.05; **p-value<0.01; ***p-value<0.001; ****p-value<0.0001.

### Extracellular GSH supplies amino acids to promote cancer cell growth and survival in cystine-free environments

The uptake of extracellular cystine is suggested to be a major source of cysteine for tumors (Fig. 2a). Interestingly, in contrast to GSH, cystine levels were lower in the TIF and plasma compared cell culture media (Fig. 2b and Extended Data Fig. 1b). GSH is a cysteine-containing tripeptide that can be broken down by GGT enzymes into cysteinylglycine and glutamate. After which cysteinylglycine can further be processed to produce intracellular cysteine. We sought to determine whether GSH or its catabolic product cysteinylglycine could replace cystine as a source of cysteine for cancer cells. We found that supplementation with GSH, at a concentration within the range found in the interstitial fluid of breast tumors *in vivo*, or cysteinylglycine rescued cell numbers in cystine-free conditions in models of breast, pancreatic, lung, and kidney cancer (Fig. 2c and Extended Data Fig. 2a). Further analysis of breast cancer cells found that GSH or cysteinylglycine rescued both survival and proliferation under cystine-free condition (Fig. 2d-2e). Cysteine deprivation is linked to ferroptosis, a non-apoptotic form of cell death that involves lipid peroxidation^52^. Interestingly, supplementation with ferrostatin-1^53^, a radical-trapping antioxidant and potent inhibitor of ferroptosis, rescued survival but not proliferation under cystine-free conditions (Extended Data Fig. 2b,c). Unlike GSH, supplementation with ferrostatin-1 does not supply cysteine, suggesting that only preventing lipid peroxidation following cystine deprivation is not sufficient to sustain cancer cells. Indeed, treatment of buthionine sulfoximine (BSO), a specific inhibitor of GSH synthesis, depleted intracellular GSH levels but did not prevent extracellular GSH from rescuing cells (Extended Data Fig. 2d,e). Next, metabolite levels were compared across control (cystine-supplemented), cystine-free, and cystine-free/GSH-supplemented media conditions. Supplementation with GSH resulted in a time-dependent accumulation of extracellular cysteinylglycine (Fig. 2f), suggesting GSH catabolism by cancer cells. While levels of intracellular cysteinylglycine and cysteine were not elevated (Fig. 2g-2h), GSH supplementation rescued the levels of downstream cysteine-related products (Fig. 2i-2k and Extended Data Fig. 2f,i). Together, these findings suggest that extracellular GSH is broken down to fuel the intracellular metabolism of cysteine by cancer cells.

**Figure 2.**
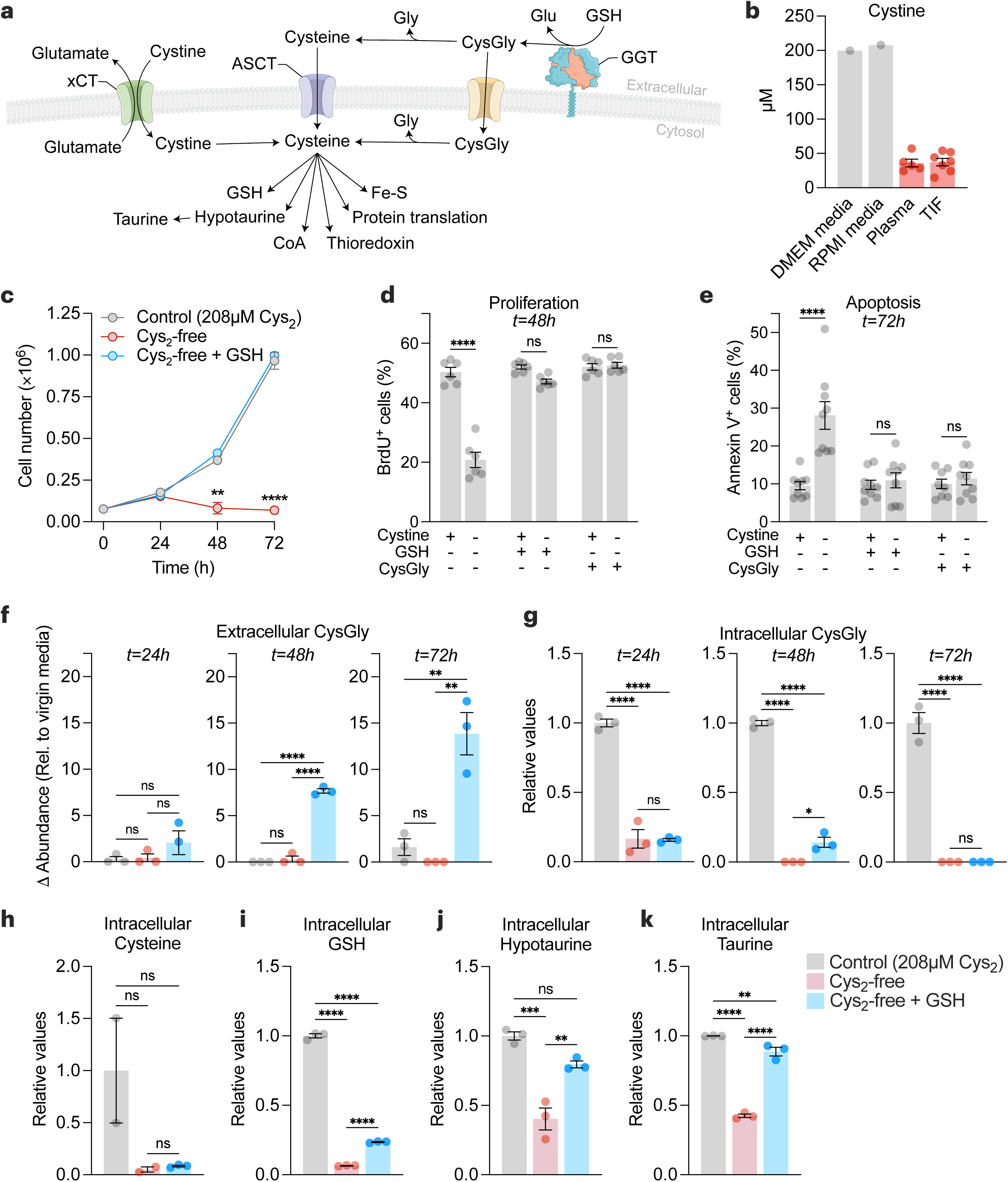
Extracellular GSH supplies amino acids to promote cancer cell growth and survival in cystine-free environments. **a**, Schematic of the different mechanisms of cysteine acquisition and utilization. **b**, Concentration of cystine in cell culture media formulations (grey bars), and in tumor interstitial fluid (n=7) and matched plasma (n=5) from Balb/c mice bearing 4T1 orthotopic allografts (red bars). **c-e**, HCC-1806 breast cancer cells were grown in control (208 µM cystine; Cys_2_), cystine-free (Cys_2_-free), cystine-free/GSH-supplemented (750 µM) or cystine-free/CysGly-supplemented (750 µM) medium and cell numbers **(c)**, percentages of proliferative (BrdU+) **(d)** and apoptotic (Annexin V+) **(e)** cells was determined at indicated timepoints. Statistical significance was analyzed by two-way ANOVA followed by Tukey’s multiple comparisons test (n=4 independent experiments) for **(c)**, two-way ANOVA followed by Šídák’s multiple comparisons test (n=3 independent experiments) for **(d-e)**. **f-g**, Levels of extracellular CysGly in media **(f)** and intracellular CysGly in HCC-1806 breast cancer cells **(g)** grown in indicated media and at indicated time points. **h-k**, Levels of intracellular cysteine **(h)**, GSH **(i)**, hypotaurine **(j)**, and taurine **(k)** in indicated media and at indicated time points. Statistical significance was assessed by one-way ANOVA followed by Tukey’s multiple comparisons test. Data is representative of an experiment with 3 biological replicates and is represented as mean ± s.e.m., *p-value<0.05; **p-value<0.01; ***p-value<0.001; ****p-value<0.0001; ns, not significant.

### GGT1 is sufficient to provide an environment permissive of cell survival under cystine-free conditions

Several enzymes possess gamma-glutamyl transferase activity^54^, with GGT1 being the most catalytically active isoform^55^. Reducing *GGT1* expression failed to prevent GSH-dependent rescue in cystine-free conditions (Extended Data Fig. 3 and 4), suggesting that GGT1 alone was not necessary for GSH catabolism and cysteine supply to cancer cells. Indeed, cancer cells express multiple GGT isoforms (Extended Data Fig. 5a,d), and none of these isoforms display dependency in cancer cell lines screening (Extended Data Fig. 5e). Alternatively, we hypothesized that GGT1 was potentially sufficient to support the breakdown of GSH in the extracellular environment. Overexpression of GGT1 in cells (i.e., GGT1^+^)^56^ resulted in higher GGT1 protein levels (Fig. 3a) and GGT activity (Fig. 3b and Extended Data Fig. 6a) and no differences in growth in cystine-supplemented conditions (Extended Data Fig. 6b). Lower levels of GSH were required to rescue GGT1^+^ cells in cystine-free conditions compared to control cells (Fig. 3c-3d), suggesting that catabolism by GGT is potentially a rate-limiting step in GSH-dependent rescue of cancer cells. We hypothesized that GSH catabolism by cells with high GGT activity could support surrounding cells in a paracrine fashion. To test this, we co-cultured GGT1^+^ cells with wild-type (WT) cells using a transwell assay (Fig. 3e). Even though GSH levels were below the threshold for rescuing WT cells in cystine-free conditions, co-culturing with GGT1^+^ cells permitted a complete rescue of WT cells (Fig. 3f). These findings demonstrate that GGT activity is sufficient to support GSH catabolism and survival of surrounding cells in cystine-depleted conditions. Furthermore, this suggests that non-tumorigenic tissues or cells in the tumor microenvironment with high GGT activity could potentially drive tumor growth and progression by catabolizing GSH and supplying amino acids in a paracrine (or endocrine) manner.

**Figure 3.**
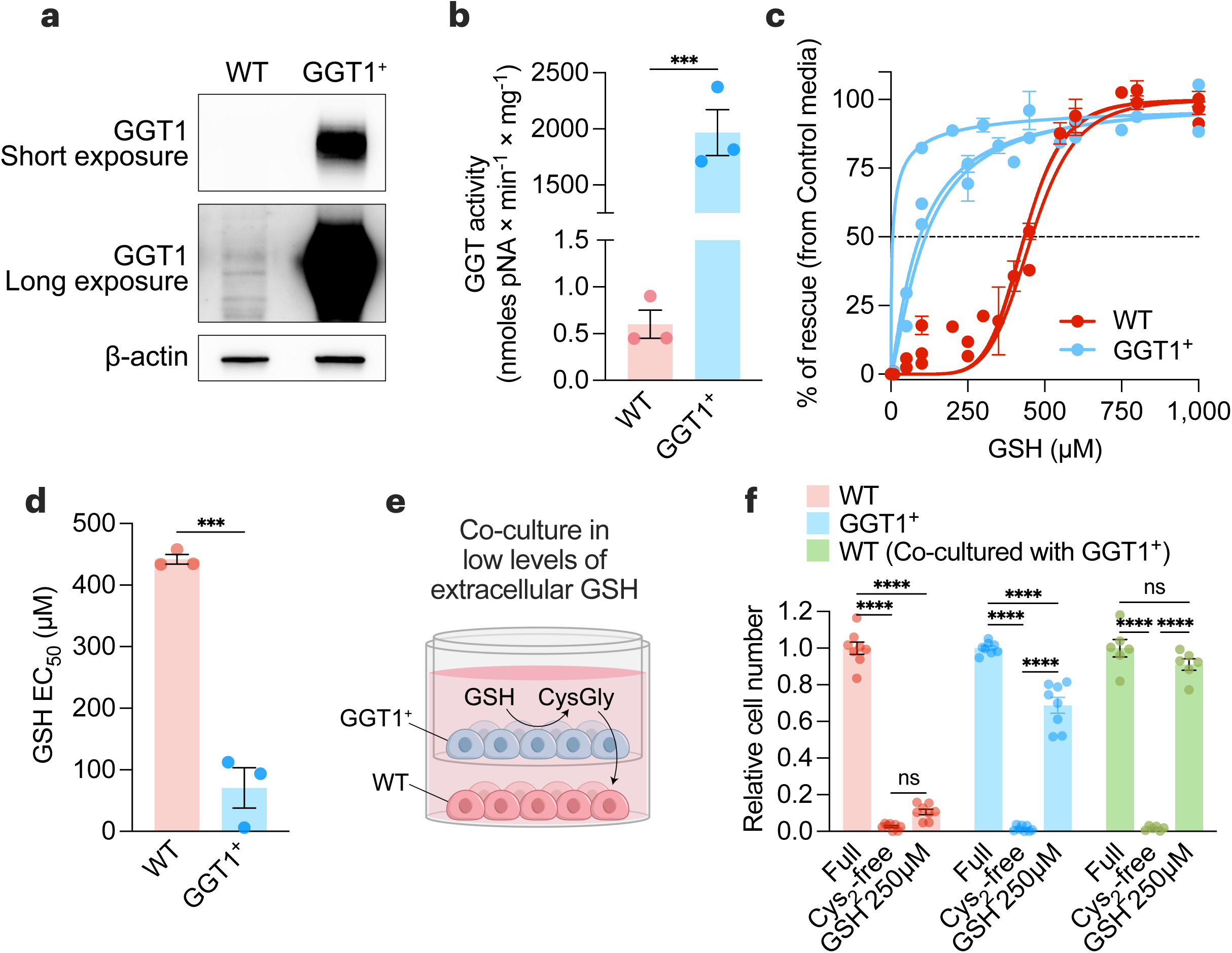
GGT1 is sufficient to provide an environment permissive of cell survival under cystine-free conditions. **a-b**, Immunoblot analysis of human GGT1 **(a)** and GGT activity **(b)** in wild-type (WT) and GGT1-overexpressing (GGT1^+^) PC3 prostate cancer cells. Statistical significance was assessed by unpaired two-tailed t-test (n=3 with two independent experiments). **c,** WT and GGT1^+^ PC3 cells were grown in control media (208 µM cystine) or cystine-free media supplemented with indicated GSH concentrations. After 72 hours, cell numbers were quantified, and the percentage of rescue ((cell numbers in indicated media – cell numbers in cystine-free media)/(cell numbers in control media) was determined (n=3 independent experiments). **d**, Half-maximal effective concentration (EC_50_) of GSH for WT and GGT1^+^ cells calculated from **(c)**. Statistical significance was assessed by an unpaired two-tailed t-test. **e,** Schematic of non-contact co-culture experiments using 0.4 μm PET membrane transwell inserts in media containing low concentrations of GSH (250 µM), which were insufficient to rescue the growth of WT cells in cystine-depleted conditions. **f,** Relative cell numbers of WT, GGT1^+^, or WT cells co-cultured with GGT1^+^ cells in control, cystine-depleted, or cystine-depleted/GSH-supplemented (250 µM) conditions. Statistical significance was evaluated by two-way ANOVA followed by Tukey’s multiple comparisons test (n=3 independent experiments). Data represented as mean ± s.e.m., ***p-value<0.001; ****p-value<0.0001; ns, not significant.

### Utilization of GSH as a cysteine source drives drug resistance in cancer cells

The metabolic environment can influence the sensitivity of cancer cells to anti-cancer compounds ^57–59^. To unbiasedly interrogate the impact on cancer cells of shifting from cystine-driven to GSH-driven cysteine supply, we performed a multifunctional approach to pharmacological screening (MAPS) with a library of 240 metabolic inhibitors (Fig. 4a-4b). We found that when cancer cells rely on GSH for cysteine acquisition, they are more sensitive to GGsTop^60^, a putative inhibitor of GGT activity (Fig. 4b-4c). Conversely, cancer cells were more resistant to inhibitors of cystine uptake (i.e., erastin) and thioredoxin pathway (i.e., auranofin, aurothioglucose, and PX-12) (Fig. 4b-4d), which mediates the reduction of cystine to cysteine following its import^61,62^. The increased resistance suggests that when cancer cells utilize GSH as a cysteine source, there is less reliance on reported anti-cancer targets (i.e., xCT and TXNRD1). In addition, the higher abundance of GSH compared to cystine in the tumor microenvironment (Fig. 1f and Fig. 2b) argues against the feasibility of targeting the proteins involved in cystine uptake and utilization for cancer therapy. Importantly, commonly used cell culture media (e.g., DMEM, RPMI, F12) contain supraphysiological levels of cystine but lack GSH and cysteinylglycine (Fig. 1f and Fig. 2b), overestimating the role of xCT while undervaluing the role of GGT in cell culture models.

**Figure 4.**
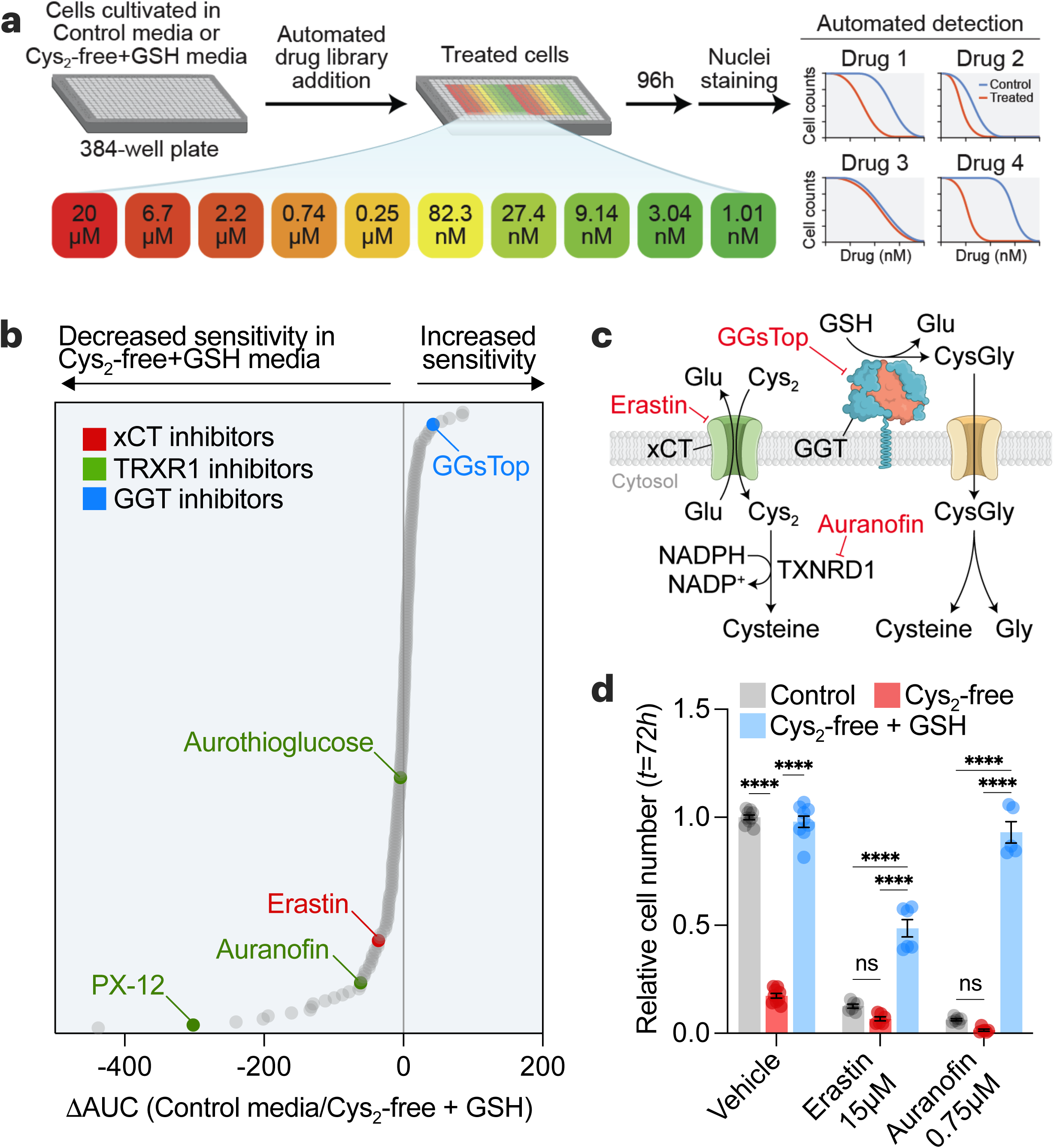
Utilization of GSH as a cysteine source drives drug resistance in cancer cells. **a**, Schematic of multifunctional approach to pharmacologic screening (MAPS). HCC1806 cells in control (208 µM cystine) or cysteine-free/GSH-supplemented (500 µM) media were treated with libraries of drugs, each arrayed at 10 dose points (20 µM – 1 nM). After 96 hours, cell numbers were determined, and dose-response curves were generated for each drug. **b,** MAPS results showing each drug ranked by the difference in the area under the curve (ΔAUC) obtained for each drug curve in the conditions. **c,** Schematic showing the targets of selected hits from **(b)**. **d,** HCC1806 cells grown in 6-well plates and treated with erastin and auranofin for 72 hours in different media conditions and relative cell numbers were determined. Statistical significance was evaluated by two-way ANOVA followed by Tukey’s multiple comparisons test (n=3 independent experiments). Data represented as mean ± s.e.m., ***p-value<0.001; ****p-value<0.0001; ns, not significant. Cys_2_, cystine; CysGly, cysteinylglycine; GGT, γ-glutamyl-transpeptidase; Glu, glutamate; GSH, glutathione; Gly, glycine; NADPH, reduced nicotinamide adenine dinucleotide phosphate; NADP+, oxidized nicotinamide adenine dinucleotide phosphate.

### GSH catabolism is necessary to support cysteine supply and tumor growth

High-throughput screening of cancer cells identified a GGT inhibitor (i.e., GGsTop) with increased sensitivity when GSH was provided as a cysteine source (Fig. 4b-4c). To further explore this, we examined additional GGT inhibitors (i.e., acivicin, OU749)^63^ and found GGsTop to be the most potent at blocking GGT activity (Fig. 5a-5b, Extended Data Fig. 7a). Notably, the increased sensitivity of GGsTop was reversed when cells were provided with cysteinylglycine, the product of GGT-mediated GSH catabolism (Fig. 5c). The kidney is suggested to have the highest levels of GGT1^64^. Indeed, the kidney showed the highest activity of GGTs (Extended Data Fig. 7b,c,d), which was suppressed following the treatment of animals with GGsTop (Fig. 5d). Blocking GGT activity slowed tumor growth without any overt toxicities to animals (Fig. 5e-5h, Extended Data Fig. 7e,f). Mechanistically, GGsTop treatment caused an accumulation of serum GSH and a depletion of cysteine and cystine in the serum and tumors (Fig. 5i-5l). Finally, the impaired tumor growth incurred by GGsTop treatment was rescued by supplementation with a cell-permeable source of cysteine (i.e., n-acetyl-cysteine) (Fig. 5m). Together, these findings suggest that GGT activity maintains GSH catabolism to supply cysteine and support tumor growth. Furthermore, the data indicate that blocking GGT is a potential therapeutic strategy for patients with cancer.

**Figure 5.**
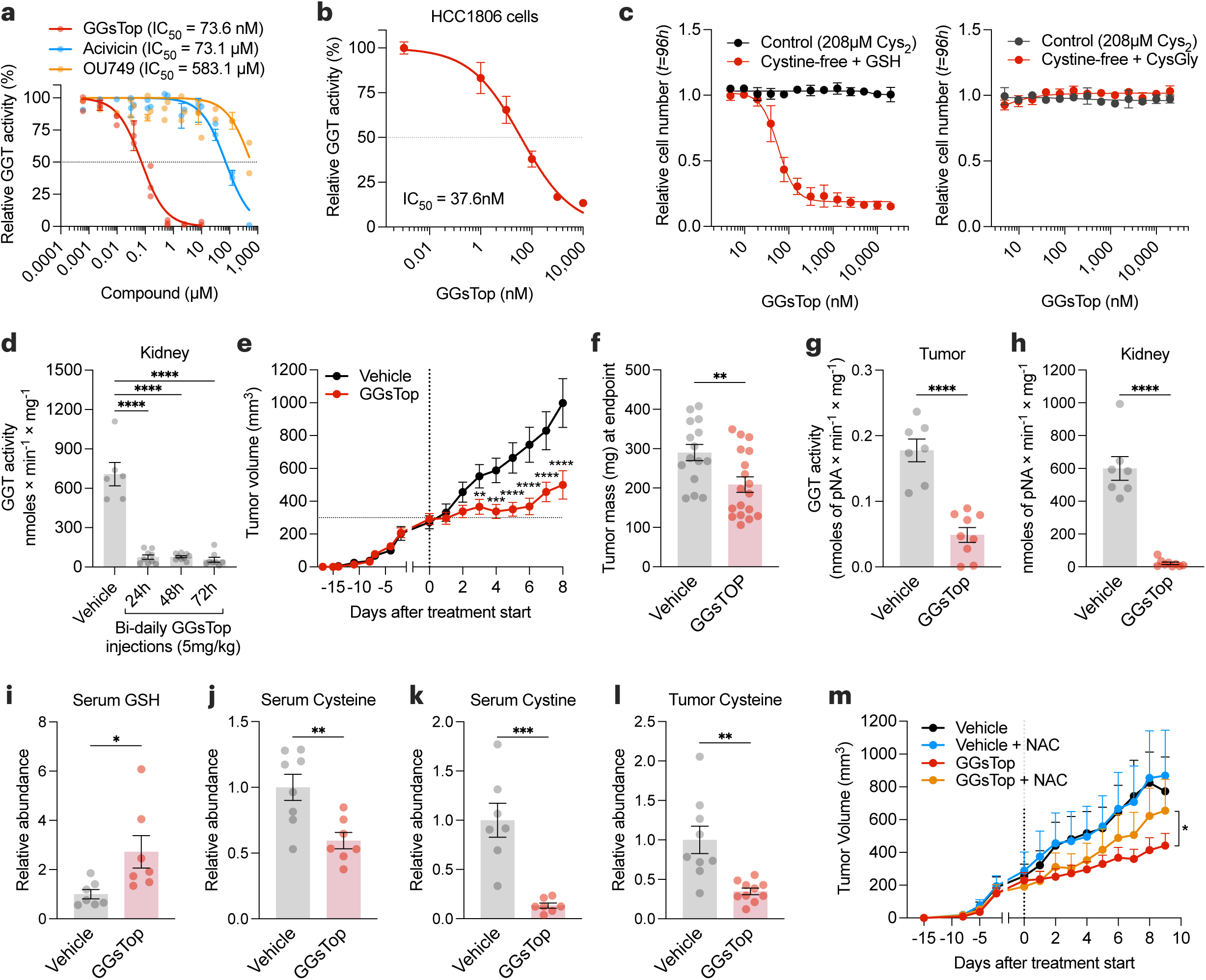
GSH catabolism is necessary to support cysteine supply and tumor growth. **a**, Mouse kidney extracts were assayed for GGT activity in the presence of GGT inhibitors. **b**, HCC-1806 cells were grown in a control medium with indicated doses of GGsTop for 4 hours, and GGT activity was determined. **c**, Relative cell number of HCC-1806 cells treated with GGsTop for 96 hours in control (208 µM cystine) or cystine-free/GSH-(left) or CysGly-supplemented (right). **d**, GGT activity in the kidney extracts of C57BL/6 mice treated intraperitoneally with vehicle (n=6 mice) or with 5 mg/kg of GGsTop every 12 hours for 1 (n=9), 2 (n=9), or 3 days (n=8). Statistical significance was evaluated by one-way ANOVA followed by Dunnett’s multiple comparisons test. **e,** Volume of orthotopically implanted HCC-1806 cell xenografts in mice treated intraperitoneally with vehicle (sterile saline) (n=15) or 5 mg/kg GGsTop (n=18) every 12 hours for 8 days. Statistical significance was assessed by two-way ANOVA followed by Tukey’s multiple comparisons test. **f-h**, Tumor mass **(f),** and GGT activity in tumor **(g)** and kidney **(h)** of mice from **(e)** at the endpoint. **i-k**, Serum levels of GSH (NEM-GSH) **(i)**, cysteine (NEM-cysteine) **(j)** and cystine **(k)** measured by LCMS. **l,** Tumor levels of cysteine (NEM-cysteine) measured by LCMS. Statistical significance in **(f-l)** was assessed by unpaired two-tailed t test. **m,** Volume of orthotopically implanted HCC-1806 cell xenografts in mice treated 5 mg/kg GGsTop every 12 hours while being supplemented with n-acetyl-cysteine (NAC, 30 mM) in their drinking water (Vehicle, n=5 mice; GGsTop, n=8; Vehicle with NAC, n=7, GGsTop with NAC, n=7). Statistical significance was assessed by two-way ANOVA. Data represented as mean ± s.e.m., *p-value<0.05; **p-value<0.01; ***p-value<0.001; ****p-value<0.0001; ns, not significant.

## Discussion

A prevailing dogma is that a major fate of cysteine is its incorporation into the antioxidant GSH. Here, we suggest that the opposite is true, that extracellular GSH is a precursor to intracellular cysteine. Additionally, we suggest that the downstream fate of cysteine in tumors is not necessarily GSH. Notably, the affinity of cysteine for cysteinyl-tRNA synthetase is orders of magnitude higher than its affinity for the rate-limiting enzyme in GSH synthesis (GCLC)^65–67^. This emphasizes that rather than synthesize GSH, cysteine has other downstream fates of greater importance, such as protein translation or cysteine polysulfidation^68^.

Our findings demonstrate that GSH catabolism by GGTs supports cancer cell growth and survival. An outstanding question is whether GSH catabolism occurs primarily by tumors or by non-tumorigenic tissue (e.g., kidney). Further, less is known whether a specific subtype of cancer relies more on GSH catabolism. Indeed, GGT1 has been studied in the context of kidney cancer, with pro- and anti-tumor effects being attributed^69–71^. Even less is known regarding its potential role in tumor metastasis and the potential side-effects of GSH catabolism within specific tissues (e.g., release of glutamate in brain metastasis). Numerous additional questions remain regarding the catabolism of metabolites and its impact on tissue homeostasis in disease.

Targeting the supply of amino acids, including cysteine, to tumors is an emerging area of therapeutic research. Significant attention has been placed on blocking cystine uptake^17,72–76^ or cysteine production from methionine via the transsulfuration pathway^9,77–79^. We examined the importance of a non-intuitive source of cysteine (i.e., GSH catabolism by GGTs). We found that blocking GGT activity slows tumor growth by depriving them of intracellular cysteine. The therapeutic window for inhibiting GGT as a cancer therapy is potentially large, as sustained inhibition of GGT activity did not elicit any overt toxicities. Degrading extracellular GSH, in a similar approach to degrading extracellular cystine/cysteine^18^, is an additional potential approach. Beyond a target for therapeutic intervention, monitoring GSH catabolism holds value as a cancer diagnostic. Indeed, serum GGT activity is a routine clinical measurement and is a risk factor for cancer^80^. Further research is required to fully elucidate the therapeutic potential surrounding GSH catabolism and the supply of amino acids to cancers.

## Methods

### Animal studies

All animal studies were performed according to protocols approved by the University Committee on Animal Resources at the University of Rochester Medical Center. *Gclc*^f/f^ mice^48^ were crossed with the MMTV-PYMT (Jackson Lab, 022974)^81^ and Rosa26-CreERT2 (Jackson Labs, 008463)^49^ mouse strains. All animals were aged for at least 12 weeks before being used in their respective experiments. To induce genetic deletion, female mice were treated with intraperitoneal (i.p.) tamoxifen (Sigma Aldrich, T5648) at 160 mg/kg once daily for five days. Orthotopic tumor allografts and xenografts occurred in females of the athymic nude NU/J (Jackson Labs, 002019) mouse strain. Allografts occurred by cell injection into the right fourth mammary fat pad or hind flank with 1×10^6^ HCC-1806 cancer cells in 100 µL sterile saline with a 29-gauge 1 mL insulin syringe. In all cases, animals only received one cell injection in one location during experimentation. Tumor volume was assessed by caliper beginning when tumors were palpable with two measurements on each tumor at the longest and shortest sides.

Tumor volume was calculated with the mathematical assumption of an oblate sphere 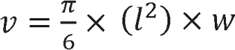 where volume *v* is calculated with the longest side always represented by *l* and the shortest side always represented by *w*. Treatment began only after the average volume of the treatment group was assessed to be approximately 300 mm^3^. Mice were treated with drugs either by i.p. injection or drinking water. GGsTop (Tocris 4452; WuXi 926281-37-0; MCE HY-108467) was diluted in sterile saline and administered via i.p. injection at indicated concentrations. N-acetyl cysteine (Sigma, A7250-100G) was diluted in drinking water at a concentration of 30 mM, and pH was adjusted to 7.00. Mice were euthanized at humane endpoints (i.e., when the tumor size exceeded a total of 20 mm in length, when ulceration occurred, or when weight loss exceeded 20%). Tumors and tissues were snap-frozen on dry ice and stored at -80 °C or placed in 10% neutral buffered formalin (Fisher Scientific 22-110-761) for 24 hours and then stored in 100% methanol (Fisher Scientific A412P-4).

### Cell culture

Cell lines were maintained in RPMI 1640 (Thermo Fisher 11875119) with 5% fetal bovine serum (FBS; Sigma SH30396.03) and 1% penicillin and streptomycin (Thermo Fisher 15070063). Cell lines were obtained from ATCC. PC3 wildtype and PC3 GGT overexpression cell lines were a gift from Dr. Marie Hannigan (Univ. of Oklahoma). For experiments involving altered media, cells were seeded in RPMI 1640 with 5% fetal bovine serum and 1% penicillin and streptomycin. RPMI 1640 medium without L-glutamine, L-cysteine, L-cystine, and L-methionine was used as the base medium for altered media (MP Biomedicals 1646454). L-glutamine (Fisher Scientific 25030081) and L-methionine (Sigma-Aldrich M5308) were added to each condition. L-Cystine (Sigma-Aldrich C7602), GSH (Sigma-Aldrich G4251), or cysteinylglycine (BACHEM 4002969) at indicated concentrations. GGsTop (Tocris 4452; WuXi 926281-37-0; MCE HY-108467), acivicin (MCE HY-W016586), OU749 (Cayman 13804), auranofin (MCE HY-B1123), erastin (MCE HY-15763) were diluted in dimethyl sulfoxide (DMSO, Sigma-Aldrich 472301) at 10 mM and added at indicated concentrations. For experiments in 6-well plates, 50,000 cells were seeded, and after 24 hours, the medium was aspirated and washed once with PBS, and the indicated medium was added. For experiments in 384-well plates, 500 cells were seeded per well, and after 24 hours, the medium was switched using a series of 4 aspiration/wash steps with the desired final medium. Co-culture experiments with PC3 WT and GGT1^+^ cell lines were performed using 6-well (24 mm) PET inserts with 0.4 µm pores (VWR 76313-902 or Corning 354570). For these experiments, cells were seeded at the same density (12,992 cells/cm^2^) in both the bottom well and the insert. For experiments in 6-well plates measuring relative cell numbers at indicated time points, the medium was aspirated, washed with PBS, and 500 µL of 0.25% trypsin (Thermo Scientific 25200056) was added to wells for 20 minutes. Next, 500 µL of RPMI 1640 with 5% fetal bovine serum and 1% penicillin and streptomycin was added, cells were resuspended, and 100 µL was added to 6.9 mL of Beckman Coulter Isoton II diluent (Fisher Scientific NC2470899) and counted using Beckman Coulter Z1 Coulter Cell Particle Counter. For experiments in 384-well plates, wells were aspirated, washed with PBS, and a fixative/staining solution containing 3.7% formaldehyde (Fisher Scientific F75P-1GAL) and 5 µg/mL Hoechst 33342, Trihydrochloride, Trihydrate (Invitrogen H1399) was added. After 30 minutes, wells were aspirated, PBS was added, and plates were sealed with adhesive foil (VWR 60941-124). Plates were imaged using the CellInsight CX5 HCS platform (Thermo), and nuclei were identified as a readout of cell counts.

### Quantification of proliferation and cell death

To quantify cell proliferation, 300,000 cells were plated in 100 mm dishes and treated as specified. At the endpoint, attached cells were pulsed with 10 mM bromodeoxyuridine (BrdU, Sigma-Aldrich B9285) for 30 minutes in their culture medium. Labeled cells were then detached by trypsinization, pelleted, washed once with PBS, resuspended in 300 µL of ice-cold PBS, and fixed by adding 700 µL of ice-cold absolute ethanol while gently vortexing. Fixed cells were stored at -20°C. For labeling, the fixed cells were thawed and washed for 5 minutes with 1 mL of 1% bovine serum albumin (BSA, Sigma-Aldrich A7906) in PBS. The cells were then centrifuged at 10,000×g for 2 minutes and incubated with 1 mL of 2 M HCl and 0.5% Triton X-100 (Sigma-Aldrich 93443) for 30 minutes with gentle agitation. Then, the cells were centrifuged and washed once with 1 mL of 0.1 M Borax (Sigma-Aldrich 71997), followed by another wash with 1% BSA in PBS. After centrifugation and thorough removal of the supernatant, the pellets were incubated for 1 hour in 60 µL of staining solution (1% BSA and 0.5% Tween-20 in PBS) containing 3 µL of anti-BrdU-FITC monoclonal antibody (Biolegend 364104, clone 3D4). The samples were subsequently washed with 1 mL of 1% BSA and 0.5% Tween-20 in PBS, centrifuged, and the pellets were resuspended in 1 mL of PI Solution (20 µg/mL propidium iodide (Sigma-Aldrich P4170), 0.5 mg/mL RNase A (Sigma-Aldrich R5503), and 0.1% Triton X-100 in PBS). The cells were incubated overnight (approximately 16 hours), and then >10,000 events were analyzed by flow cytometry using a BD Accuri™ C6 Plus Flow Cytometer (BD Biosciences) and FCS Express 7 Research (DeNovo Software).

For quantifying cell death, 50,000 HCC-1806 cells were seeded into 6-well plates and treated as indicated. At the endpoint, cells were collected by trypsinization (the culture media and PBS washes were collected and readded to each sample) and washed once with assay buffer (10 mM HEPES, 140 mM NaCl, 3.3 mM CaCl2, pH 7.4). Then, the pellets were stained for 20 minutes in 100 µL of assay buffer containing 5 µg/mL propidium iodide (Sigma-Aldrich P4170) and 3 µL of Annexin V-FITC (Biolegend 0640906). Then, cells were diluted with 0.5-1 mL of assay buffer and >10,000 events were analyzed by flow cytometry using a BD Accuri™ C6 Plus Flow Cytometer (BD Biosciences) and FCS Express 7 Research (DeNovo Software).

### Immunoblot assays

Cell lysates from PC3 wildtype and PC3 GGT overexpression cells were obtained by placing PBS-washed cell plates on ice and scraping with RIPA buffer (Thermo Scientific 89900) containing Halt Protease & Phosphatase inhibitor cocktail (Thermo Scientific 1861280). Cell lysates were incubated for 30 minutes on ice and then centrifuged at 12,000×g for 15 minutes at 4 °C. The supernatant was collected and stored at -80 °C. Extracted proteins were quantified using the Pierce BCA Protein Assay Kit (Thermo Scientific 23225). 75-100µg of protein lysates were mixed with Laemmli 6x SDS sample buffer (Boston BioProducts, BP-111R) with 5% 2-mercaptoethanol (VWR Life Science M131) and then ran on 4-20% Criterion TGX pre-cast gels (Bio-Rad 5671093). Separated proteins were transferred onto Immobilon-P Transfer membranes (MilliporeSigma IPVH00010), blocked for 1 hour using 5% milk in TBS-0.5% Tween-20, and stained overnight with affinity-purified rabbit-anti-GGT1 (GGT129, described in ^82^) 1:1,000 in BSA 2.5% overnight at 4 °C. Stained membranes were washed three times with TBS-T and stained with secondary antibody donkey-anti-rabbit (Thermo Fisher NAV934V) 1:5,000 in BSA 2.5% for 1 hour. Membranes were washed 3 more times with TBS-T, and then the antibody-stained protein signal was amplified and visualized using SuperSignal (ThermoFisher 34577). Blots were imaged with a ChemiDoc MP Imaging system (Bio-Rad). Membranes were stripped for 20 min at room temperature using Blot Restore Membrane Rejuvenation Kit (Sigma 2520-M), then washed twice with TBS-T, re-blocking with BSA 5% for 30 minutes and incubated with mouse-anti-β-actin (Sigma A1978) 1:3,000 in BSA 2.5% for 4 hours at room temperature and then incubated with a sheep-anti-mouse (Thermo Fisher NA931V) secondary antibody at 1:5,000 in BSA 2.5% for 2 hours.

### Drug screening

MAPS was performed as previously described^57^. Briefly, 500 cells were seeded per well in 384-well plates, and after 24 hours, the medium was switched using a series of 4 aspiration/wash steps with the desired final medium. Compound libraries were gifts from Dr. Joan Brugge, the Ludwig Center at Harvard Medical School, and the ICCB-L. 100 nL of compounds were transferred using the JANUS MDT Workstation (Revvity) and a 384-well pin tool (V&P Scientific). After 96 hours, wells were aspirated, washed with PBS, and a fixative/staining solution containing 3.7% formaldehyde (Fisher Scientific F75P-1GAL) and 5 µg/mL Hoechst 33342, Trihydrochloride, Trihydrate (Invitrogen H1399) was added. After 30 minutes, wells were aspirated, PBS was added, and plates were sealed with adhesive foil (VWR 60941-124). Plates were imaged using the CellInsight CX5 HCS platform (Thermo), and nuclei were identified as a readout of cell counts. Data post-processing was conducted using R scripts.

### GGT activity assay

Cell cultures were washed and scraped with PBS. The use of trypsin for cell detachment is avoided to prevent proteolytic degradation of cell-surface GGT. Detached cells were collected and centrifuged 300×g for 5 minutes at room temperature. Then, pellets were resuspended in lysis buffer (Tris-HCl 100 mM, 0.5% Triton X-100, pH 8) containing Halt Protease and Phosphatase Inhibitor Cocktail (Thermo Scientific 1861280), thoroughly vortexed for 1 minute and incubated for 15 minutes on ice. Lysates were cleared by centrifugation at 12,000×g for 15 minutes at 4 °C and the supernatants were stored at -80 °C until analysis. Tissues were lysed by transferring them to pre-filled bead mill tubes (Fisher Scientific 15-340-154) containing lysis buffer (Tris-HCl 100 mM, 0.5% Triton X-100, pH 8) containing Halt Protease and Phosphatase Inhibitor Cocktail and homogenized using a bead mill (VWR) for 10 seconds. Tissue lysates were then rotated for 15 minutes at 4 °C and centrifuged at 12,000×g for 15 minutes at 4 °C. Supernatants of cells or tissues were stored at -80 °C until analysis. Extracted proteins were quantified using the Pierce BCA Protein Assay Kit. GGT reaction was carried out by using the GpNA method^83^. Briefly, cell or tissue protein lysates were assayed in transparent 96-well plates (Greiner 655101) in the presence of 1 mM of the GGT substrate L-glutamic acid γ-p-nitroanilide (GpNA, Cayman 36209) and 20 mM of the transpeptidation acceptor glycyl-glycine (Sigma-Aldrich G1002) in Tris-HCl 100 mM pH 8. GGT catalyzes the breakdown of the γ-glutamyl bond in the substrate and generates p-nitroanilide (pNA), which is monitored kinetically by an increase in absorbance at 418nm. P-nitroaniline (Sigma 185310) was used to generate a standard curve (0-40nmoles) for the calculation of GGT activity. Absorbance (418 nm) was assessed using a plate reader set at 37 °C with a reading time of one hour and 2-5 minutes intervals between reads. GGT activity was calculated by obtaining the slope of the linear range of each sample (Abs/min) and dividing by the slope of a 4-nitroaniline standard curve (Abs/nmoles pNA) and by the protein mass (mg) to determine the reaction rate of the sample (nmoles pNA × min^-1^ × mg^-1^). The optimal protein mass for the GGT assay was determined experimentally for each tissue or cell line (HCC-1806 lysates or tumor xenografts: 200 µg, PC3 cell line: 100 µg, PC3-GGT1^+^: 1 µg, kidney: 0.5 µg, spleen: 150 µg, Liver: 250 µg, seminal vesicle: 10 µg, pancreas: 2.5 µg, lungs: 2.5 µg, muscle: 200 µg, testis: 50 µg, epididymis: 10 µg, PyMT breast tumor: 50 µg, heart: 150 µg, mammary fat pad: 150 µg, brain: 80 µg, large intestine: 80 µg, brown adipose tissue: 300 µg).

### GGT histochemical assay

The histochemical staining of GGT was performed as previously described^84^, with modifications. Cells were seeded into 6-well plates and cultured in complete media. At the endpoint, cells were washed three times with saline and stained for 20 minutes at room temperature using a saline solution containing 0.2 mM glutamic acid gamma-4-methoxy-beta-naphthylamide (GMNA, Sigma-Aldrich G0141), 20 mM glycyl-glycine (Sigma-Aldrich G1002), 25 mM Tris (Sigma-Aldrich 93362), and 1.2 mM FastBlue (Sigma-Aldrich F3378). For negative controls, the staining solution included 5 mM serine (Sigma-Aldrich S4311) and 10 mM sodium borate (Sigma-Aldrich 71997). After incubation, cells were rinsed with saline and incubated with 100 mM CuSO4 (Sigma-Aldrich C1297) for 2 minutes. Following a final saline wash, 50% glycerol was added to the wells, and brightfield images were taken using an inverted microscope.

### GSH quantification

For some experiments, glutathione concentration was determined using the GSH-Glo^TM^ Glutathione Assay (Promega V6912). Briefly, 6,250 cells were seeded into 96-well white cell culture-treated plates (Falcon 353296), treated for 24 hours as indicated, and labeled following the manufacturer’s instructions. The luminescence of samples and of the glutathione standard curve was detected using a Victor multiplate reader (Perkin Elmer).

### RNA analysis

mRNA was isolated from cells using E.Z.N.A. total RNA Kit I (Omega Bio-Tek R6834-02). For gene expression analysis, 1 µg of RNA was used for cDNA synthesis using qScript cDNA Synthesis Kit (Quanta Bio 66196756). The expression of target genes was analyzed via Quantitative real-time (RT) PCR with a QuantaStudio 5 qPCR machine (Applied Biosystems, Thermo Fisher Scientific).

### Serum analysis

Mice were anesthetized with isoflurane, after which blood was collected via the retro-orbital venous sinus into BD microtainer tubes (BD 365967). Serum was isolated from the blood by centrifuging blood samples at 10,000×g for 5 minutes and stored at -80 °C. VRL Animal Health Diagnostics carried out the analysis of liver damage biomarkers.

### Isolation of plasma and TIF from 4T1 breast cancer tumor-bearing mice

4T1 cells were acquired from ATCC and cultured in RPMI supplemented with 10% FBS and 0.5% penicillin and streptomycin at 37°C in a 5% CO_2_ tissue culture incubator. 5×10^5^ cells in Matrigel were injected into the #9 mammary glands of 6-8 week-old Balb/c mice (Jackson Labs). TIF was isolated using a previously described centrifugation method^85^. Briefly, once the tumors reached approximately 1 to 1.5 cm in diameter, mice were euthanized by cervical dislocation, and tumors were dissected within 1 minute. Excised tumors were briefly rinsed with saline and blotted to remove residual fluid prior to centrifugation at 100×g for 10 min at 4°C using 10 µm nylon filters (Spectrum Labs, 148134) affixed to 50 mL conical tubes. The flow-through was collected into an Eppendorf tube and centrifuged at 845×g for 10 mins at 4°C to eliminate the residual cellular debris. Isolated TIF was flash-frozen in liquid nitrogen and stored at -80°C. Blood samples were collected by cardiac puncture into EDTA tubes and centrifuged at 845×g for 10 mins at 4°C. Plasma was in Eppendorf tubes and flash-frozen in liquid nitrogen prior to storage at -80°C.

### Metabolite analysis

Cells were seeded in 6-well plates and treated as specified. At the endpoint, cells were quickly washed with ice-cold PBS and incubated at -80 °C for 30 minutes with the extraction solvent at a ratio of 500 µL per 5×10^5^ cells. The cells were then scraped on ice, transferred to pre-chilled microcentrifuge tubes, centrifuged at 15,000×g for 20 minutes, and the supernatants collected. Conditioned media were extracted by taking 10 µL of the media supernatant and mixing it with 390µL of the extraction solvent containing NEM. Virgin media controls were generated by incubating the treatment media in cell-free 6-well plates under the same conditions and duration as the cell culture samples. Media extracts were incubated at -80 °C for at least 15 minutes before being centrifuged at 15,000×g for 20 minutes. All extracts were stored at -80 °C. The extraction solvent consisted of 80% methanol and 20% aqueous solution (pH 7.00) containing 10 mM ammonium formate (Sigma-Aldrich 70221-25G-F) and 25 mM N-ethylmaleimide (Thermo Scientific 040526.06). The final concentrations of ammonium formate and NEM in the extraction solvent were 2 mM and 5 mM, respectively. For chromatographic metabolite separation, a Vanquish ultra-performance liquid chromatography (UPLC) system was coupled to a Q Exactive HF (QE-HF) mass spectrometer equipped with heated electrospray ionization (HESI; Thermo Fisher Scientific, Waltham, MA, USA). Samples were run on an Atlantis Premier BEH Z-HILIC VanGuard FIT column, 2.5 µm, 2.1 mm x 150 mm (Waters, Milford, MA, USA). The mobile phase A was 10 mM (NH_4_)_2_CO_3_ and 0.05% NH_4_OH in H_2_O, while mobile phase B was 100% acetonitrile (ACN). The column chamber temperature was maintained at 30°C. The mobile phase parameters were set according to the following gradient: 0-13min: 80% to 20% of mobile phase B, 13-15min: 20% of mobile phase B. The ESI ionization mode was negative, and the MS scan range (m/z) was set to 65-975. The mass resolution was 120,000 and the AGC target was 3×10^6^. The capillary temperature and capillary voltage were maintained at 320 °C and 3.5 kV, respectively. 5mL of each sample was loaded for metabolite detection. The LC-MS metabolite peaks were manually integrated and identified by El-Maven (Version 0.12.0) by matching with a previously established in-house library^86^.

Frozen tumor tissue was homogenized in 80% MeOH containing 20 mM NEM using a Precellys cold tissue homogenizer (Bertin), at a ratio of 20mg tissue to 1 mL solvent. Following homogenization, samples were transferred to -80 °C for 30 min and then placed on regular ice for 30 minutes with vortexing every 10 minutes. Next, samples were centrifuged at 17,000×g for 10 minutes and 800 µL supernatant was dried down in a vacuum evaporator (Thermo).

Samples were reconstituted in 90µL of 50% acetonitrile (A955, Fisher Scientific) and transferred to glass vials for LC/MS analysis. For mice serum collection, the animals were anesthetized with isoflurane, and blood was collected via the retro-orbital venous sinus into BD microtainer tubes (BD 365967). Serum was obtained by centrifuging blood samples at 10,000×g for 5 minutes and stored at -80 °C. Then, 20 µL of serum was mixed with 2 µL of 200 mM N-ethylmaleimide (NEM, Thermo Scientific 040526.06) and extracted with 80% methanol. Next, 900 µL of serum extract was dried down in a vacuum evaporator (Thermo), reconstituted in 90 µL of 50% acetonitrile (A955, Fisher Scientific) and transferred to glass vials for LC/MS analysis. LC-MS analysis was carried out by the URMC Metabolomics Resource. Metabolite extracts were analyzed by high resolution mass spectrometry with an Orbitrap Exploris 240 (Thermo) coupled to a Vanquish Flex liquid chromatography system (Thermo). 2 µL of samples were injected on a Waters XBridge XP BEH Amide column (150 mm length × 2.1 mm id, 2.5 µm particle size) maintained at 25 °C, with a Waters XBridge XP VanGuard BEH Amide (5 mm × 2.1 mm id, 2.5 µm particle size) guard column. For positive mode acquisition, mobile phase A was 100% LC-MS grade H_2_O with 10 mM ammonium formate and 0.125% formic acid. Mobile phase B was 90% acetonitrile with 10 mM ammonium formate and 0.125% formic acid. For negative mode acquisition, mobile phase A was 100% LC-MS grade H_2_O with 10 mM ammonium acetate, 0.1% ammonium hydroxide, and 0.1, 100% B; 2□minutes, 100% B; 3□minutes, 90% B; 5□minutes, 90% B; 6□minutes, 85% B; 7□minutes, 85% B; 8□minutes, 75% B; 9□minutes, 75% B; 10□minutes, 55% B; 12□minutes, 55% B; 13□minutes, 35%, 20□minutes, 35% B; 20.1□minutes, 35% B; 20.6□minutes, 100% B; 22.2 minutes, 100% B all at a flow rate of 150□μL min^−1^, followed by 22.7□minutes, 100% B; 27.9□minutes, 100% B at a flow rate of 300□μL min^−1^, and finally 28□minutes, 100% B at flow rate of 150□μL min^−1^, for a total length of 28 minutes. The H-ESI source was operated in positive mode at spray voltage 3,500 or negative mode at spray voltage 2,500 with the following parameters: sheath gas 35 au, aux gas 7 au, sweep gas 0 au, ion transfer tube temperature 320 °C, vaporizer temperature 275 °C, mass range 70 to 800 m/z, full scan MS1 mass resolution of 120,000 FWHM, RF lens at 70%, and standard automatic gain control (AGC). LC-MS data were analyzed by El-Maven software^87^, and compounds were identified by matching to LC-MS method specific retention time values of external standards.

Quantification of metabolites in interstitial fluid and plasma by liquid chromatography mass spectrometry was performed as previously reported^6,85^. Briefly, 5 µL of biofluid samples or serial dilutions of a chemical standard library of 149 metabolites were extracted with 45 µL of a 75:25:0.1 HPLC grade acetonitrile:methanol:formic acid extraction mix that included the following labeled stable isotope internal standards:^13^C-labeled yeast extract (Cambridge Isotope Laboratory, Andover, MA, ISO1), ^2^H_9_ choline (Cambridge Isotope Laboratory, Andover, MA, DLM-549), ^13^C_4_ 3-hydroxybutyrate (Cambridge Isotope Laboratory, Andover, MA, CLM-3853), ^13^C_6_^15^N_2_ cystine (Cambridge Isotope Laboratory, Andover, MA, CNLM4244), ^13^C_3_ lactate (Sigma-Aldrich, Darmstadt, Germany, 485926), ^13^C_6_ glucose (Cambridge Isotope Laboratory, Andover, MA, CLM-1396), ^13^C_3_ serine (Cambridge Isotope Laboratory, Andover, MA, CLM-1574), ^13^C_2_ glycine (Cambridge Isotope Laboratory, Andover, MA, CLM-1017), ^13^C_5_ hypoxanthine (Cambridge Isotope Laboratory, Andover, MA, CLM8042), ^13^C_2_^15^N taurine (Cambridge Isotope Laboratory, Andover, MA, CNLM-10253), ^13^C_3_ glycerol (Cambridge Isotope Laboratory, Andover, MA, CLM-1510) and ^2^H_3_ creatinine (Cambridge Isotope Laboratory, Andover, MA, DLM-3653). Samples or chemical standards were vortexed for 10 min at 4°C and centrifuged at 21,000×g for 10 min at 4°C to pellet insoluble material. The soluble polar metabolite supernatant was then moved to sample vials for LC-MS analysis as previously described^6^. Data analysis was performed as previously described^5,85^ using Skyline^88^.

### Statistical analysis

All statistical analysis was completed using either R or GraphPad Prism 10.3.1.

## Data availability

Data supporting these findings are included within the article and its supplementary material.

Source data are provided with this paper. All data reported in this study are available from the corresponding author upon request.

## Acknowledgments

We thank Samuel McBrayer for their feedback and discussions. We thank Marie Hanigan (University of Oklahoma) for the assistance with protocols and donation of key reagents. We would also like to thank the Metabolomics Resource in the Center for Advanced Research Technologies (CART) and the Histology, Biochemistry, and Molecular Imaging (HBMI) Core at the Center for Musculoskeletal Research (CMSR) at URMC, the Proteomics/Metabolomics Core at Moffitt Cancer Center, which is funded in part by Moffitt’s Cancer Center Support Grant (P30CA076292), and University of Chicago Metabolomics Platform (RRID: SCR_022932). We also thank Joan Brugge and the Ludwig Center at Harvard Medical School for their support. This work was supported by the Wilmot Cancer Institute Predoctoral Fellowship (G.A.) the American Association for Cancer Research and Breast Cancer Research Foundation (20-20-26-HARR) (I.S.H.), Breast Cancer Coalition of Rochester (I.S.H.), American Cancer Society (RSG-23-971782-01-TBE) (I.S.H.), and NIH grants R01CA269813 (I.S.H.), R01CA269813-S1 (M.Z.), R37CA230042 (G.M.D), R24AA022057 (V.V.), R01AA028859 (Y.C.), AI150698 (J.M.).

## Author Contributions Statement

F.H., M.Z., and I.S.H. initiated the study, conceived the project, designed experiments, interpreted results, and wrote the manuscript. F.H. and M.Z. performed the experiments with assistance from E.T.T., J.C., Z.G.S., F.A., L.D.M., D.T., G.A., and S.K.B.-N.. N.P.W., Y.P.K., and G.M.D. performed metabolite analyses for in vitro experiments. B.S., J.M., C.S., and A.M. performed metabolite analysis for in vivo experiments. J.J.Z., M.E.O., and J.L.C. assisted with orthotopic allograft tumor experiments. Y.C., V.V., and B.M.T. provided reagents and expert comments.

