## Supplementary figures and images for "Catabolism of extracellular glutathione supplies amino acids to support tumor growth"

### Extended Data Figure 1

##### Extended Data 1

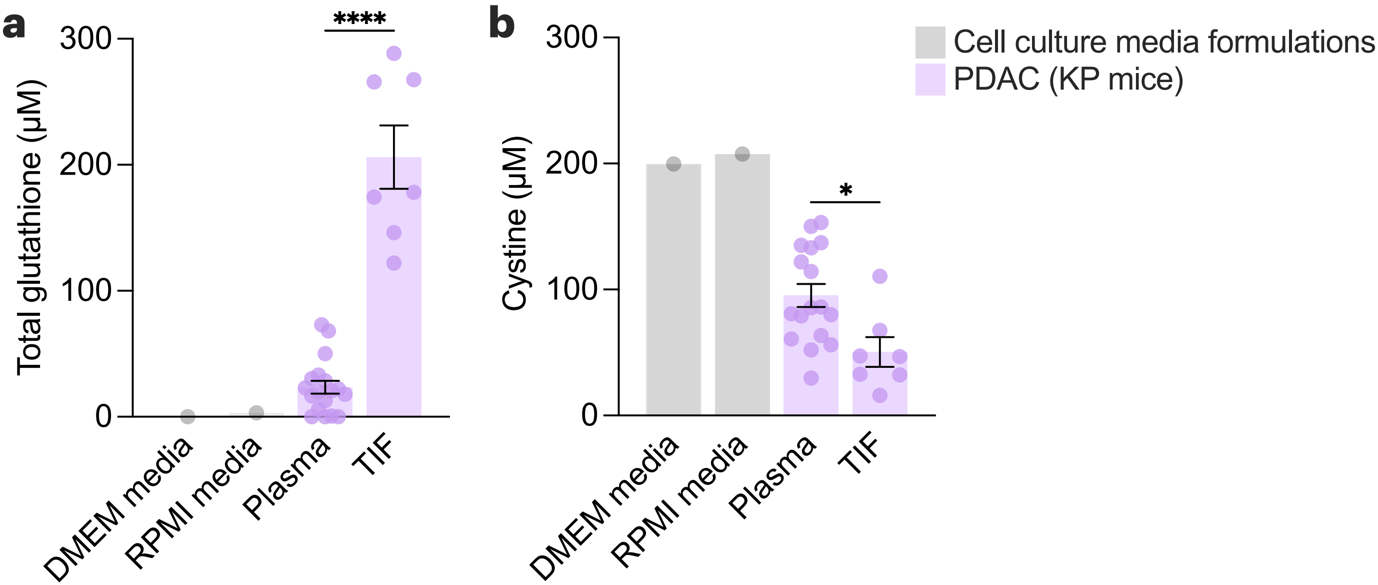

### Extended Data Figure 2

#### Extended Data 2

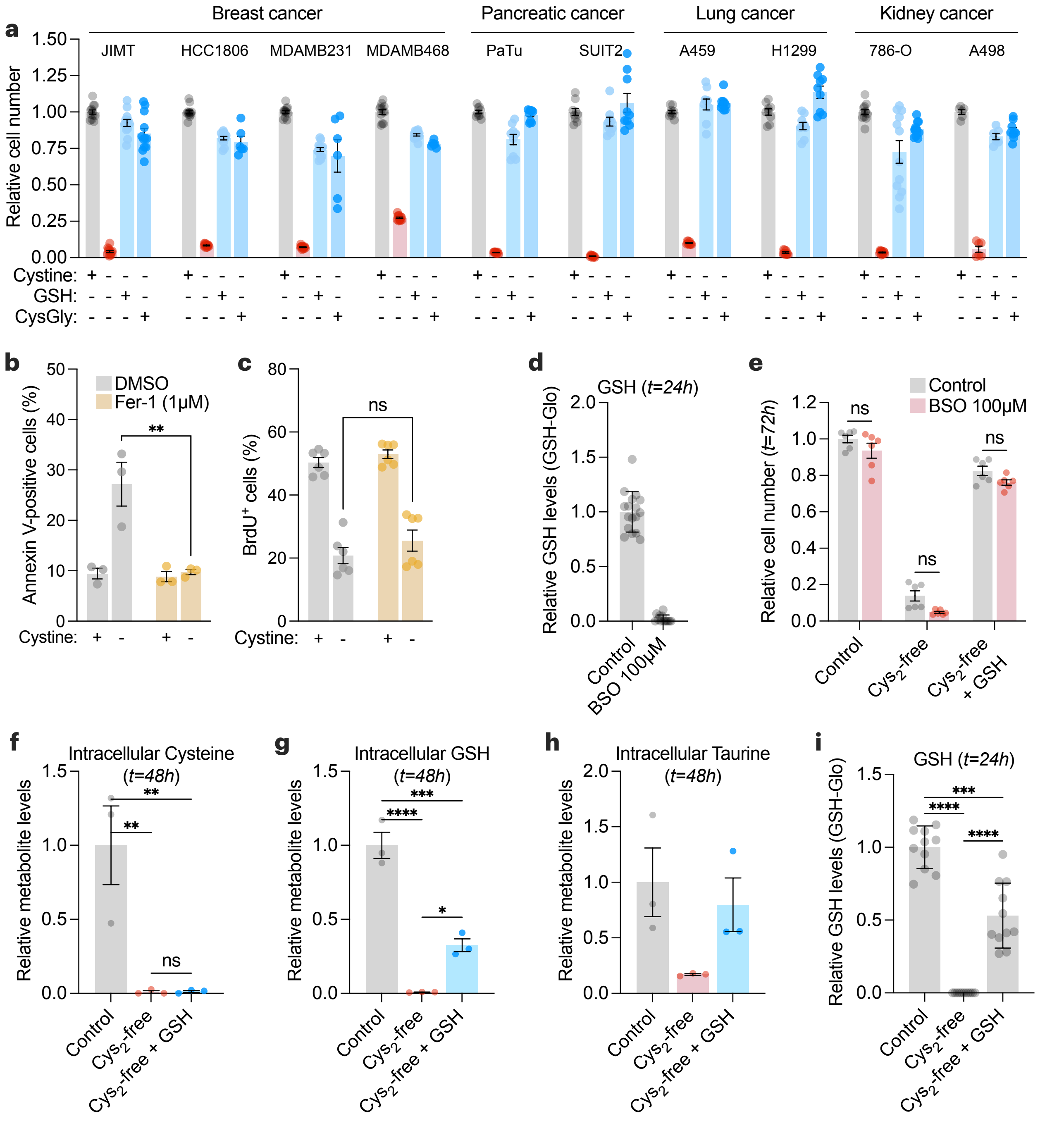

### Extended Data Figure 3

##### Extended Data 3

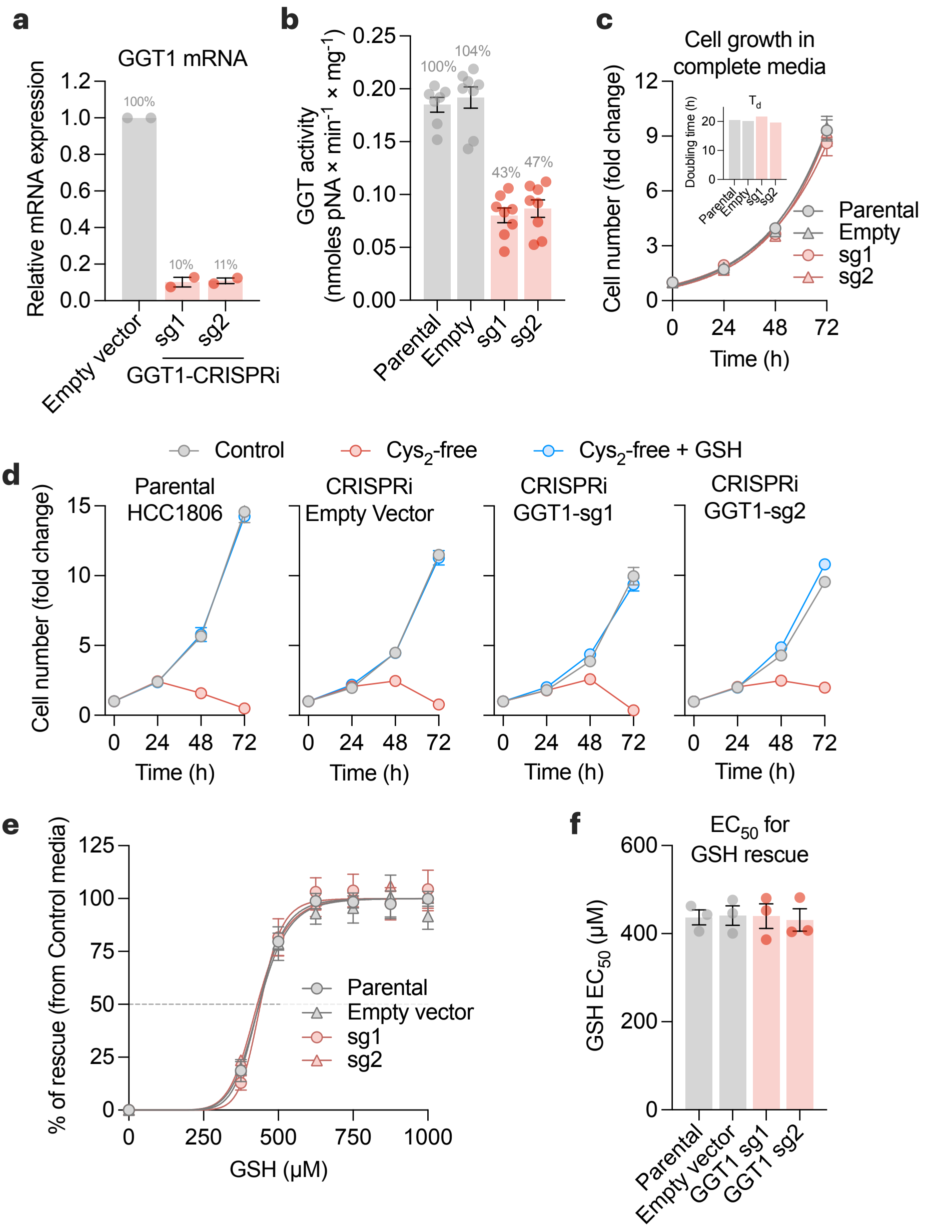

### Extended Data Figure 4

**Extended Data 4**

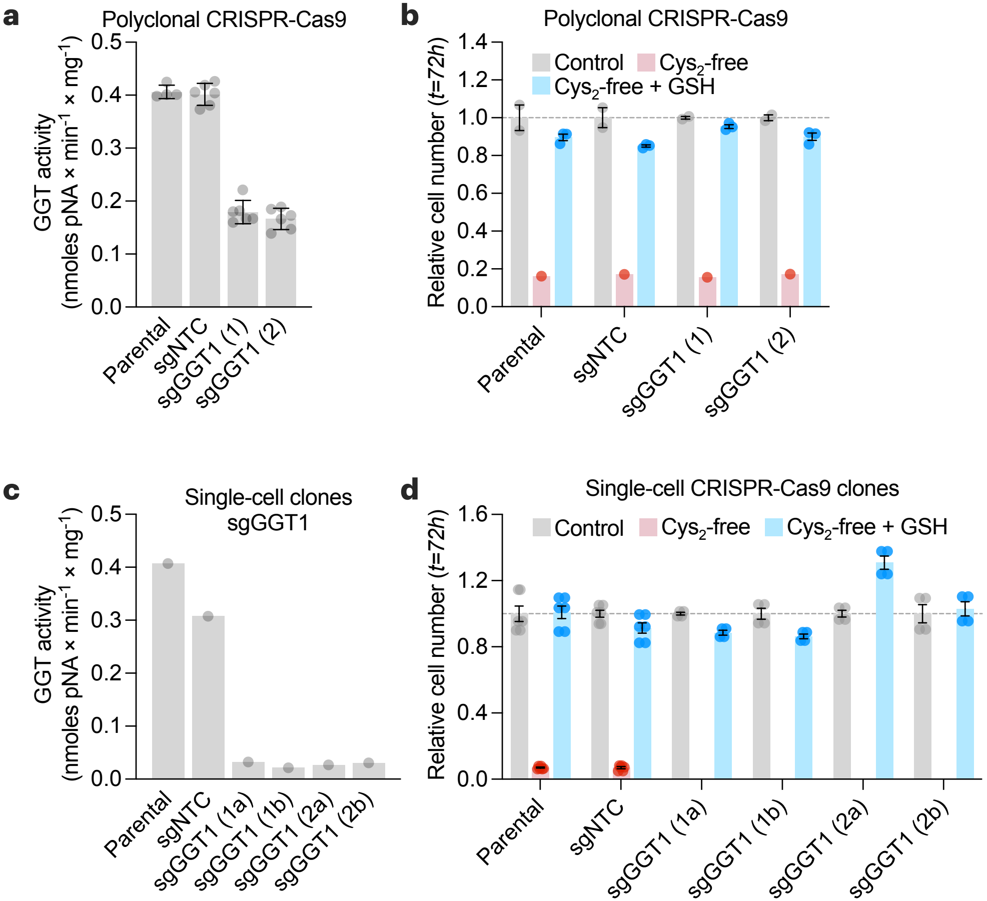

### Extended Data Figure 6

#### Extended Data 6

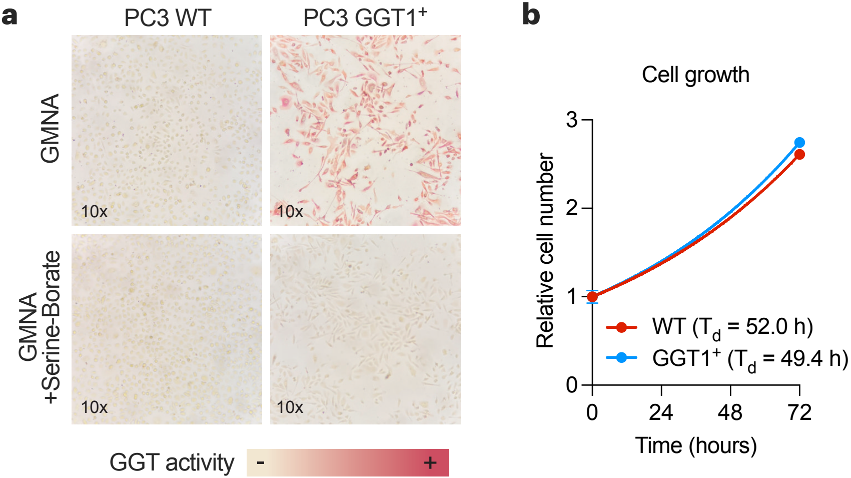

### Extended Data Figure 7

##### Extended Data 7

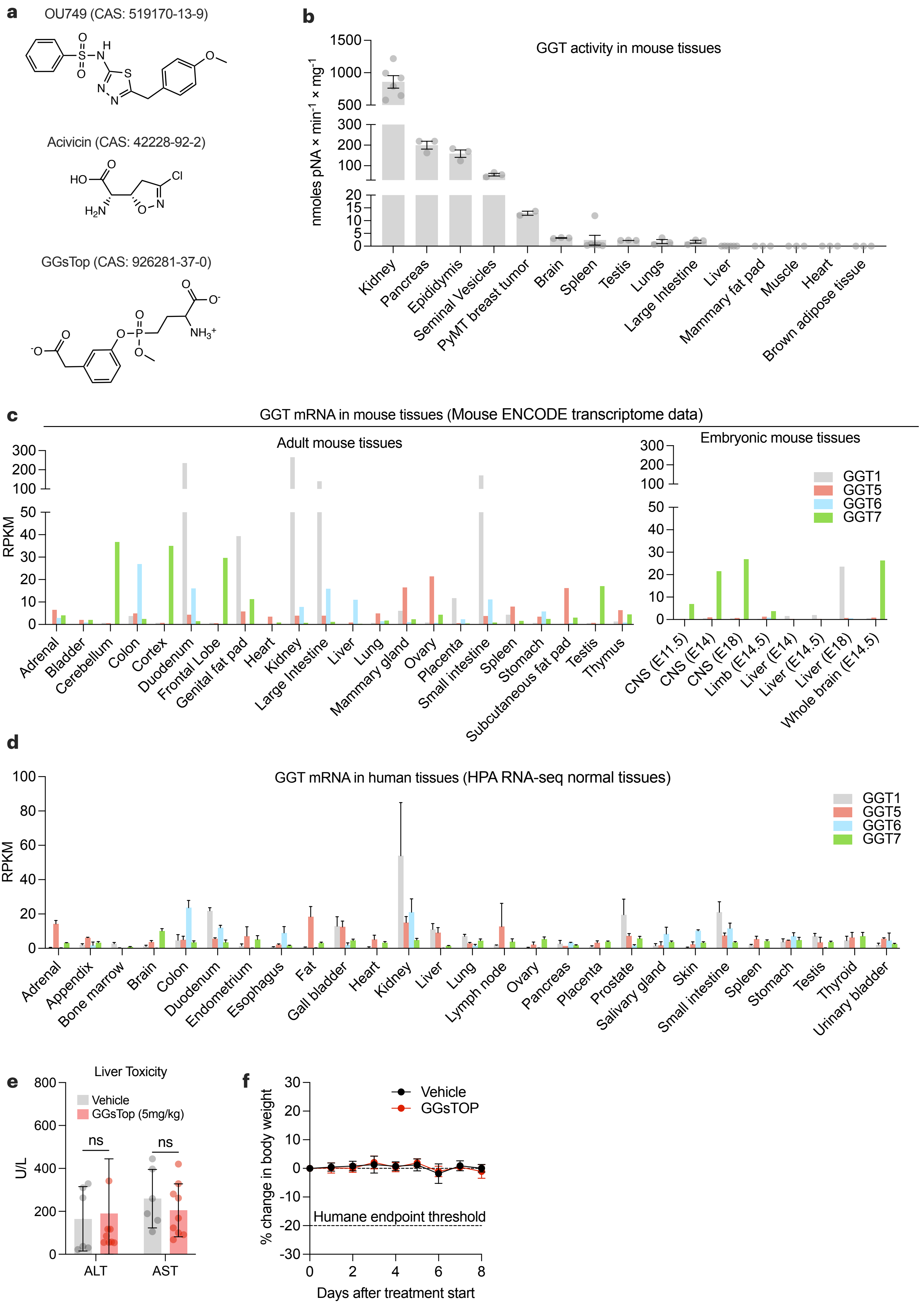
