## Extended Data Figure 5 for "Catabolism of extracellular glutathione supplies amino acids to support tumor growth"

### Extended Data 5

**a**

| | Gene | Location | NCBI code | Gene type | Homology to GGT1 (amino acid level) | Enzymatic activity | Ref |
|---|---|---|---|---|---|---|---|
| 1 | GGT1 | 22q11.23 | 2678 | Protein coding (full length) | - | Active (experimentally confirmed) | 1,2 |
| 2 | GGT5 | 22q11.23 | 2687 | Protein coding (full length) | High | Active (experimentally confirmed) | 1,2 |
| 3 | GGT6 | 17p13.2 | 124975 | Protein coding (full length) | Limited | Not charaterized, enzymatic activity possible | 2 |
| 4 | GGT7 | 20q11.22 | 2686 | Protein coding (full length) | High | Not charaterized, enzymatic activity possible | 2 |
| 5 | GGTLC1 | 20p11.21 | 92086 | Protein coding (light chain only) | | Not charaterized, enzymatic activity possible | 2 |
| 6 | GGTLC2 | 22q11.22 | 91227 | Protein coding (light chain only) | | Not charaterized, enzymatic activity possible | 2 |
| 7 | GGTLC3 | 22q11.21 | 728226 | Protein coding (light chain only) | | Not charaterized, enzymatic activity possible | 2 |
| 8 | GGT2P | 22q11.21 | 728441 | Pseudogene | | Inactive (experimentally confirmed) | 2,3 |
| 9 | GGT3P | 22q11.21 | 2679 | Pseudogene | | Inactive (predicted) | 2 |
| 10 | GGT4P | 13q11 | 643171 | Pseudogene | | Inactive (predicted) | 2 |
| 11 | GGT8P | 2p11.2 | 645367 | Pseudogene | | Inactive (predicted) | 2 |
| 12 | GGTLC4P | 22q11.23 | 729838 | Pseudogene | | Inactive (predicted) | 2 |
| 13 | GGTLC5P | 22q11.21 | 653590 | Pseudogene | | Inactive (predicted) | 2 |

References (doi): 1 (10.1016/j.ab.2011.03.026), 2 (10.1007/s00439-008-0487-7), 3(10.1089/ars.2012.4997)

**b**

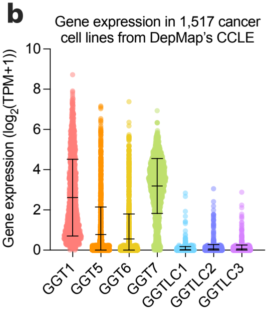


**c**

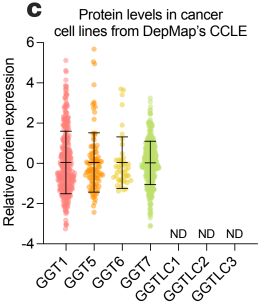


**d**

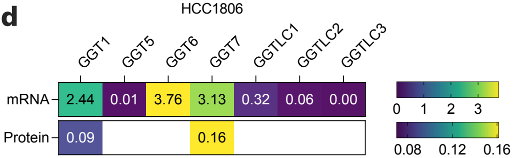


**e**

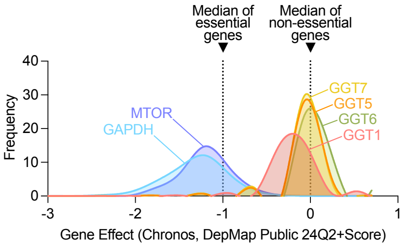
